# aaHash: recursive amino acid sequence hashing

**DOI:** 10.1101/2023.05.08.539909

**Authors:** Johnathan Wong, Parham Kazemi, Lauren Coombe, René L. Warren, Inanç Birol

## Abstract

**Motivation:** *K*-mer hashing is a common operation in many foundational bioinformatics problems. However, generic string hashing algorithms are not optimized for this application. Strings in bioinformatics use specific alphabets, a trait leveraged for nucleic acid sequences in earlier work. We note that amino acid sequences, with complexities and context that cannot be captured by generic hashing algorithms, can also benefit from a domain-specific hashing algorithm. Such a hashing algorithm can accelerate and improve the sensitivity of bioinformatics applications developed for protein sequences.

**Results:** Here, we present aaHash, a recursive hashing algorithm tailored for amino acid sequences. This algorithm utilizes multiple hash levels to represent biochemical similarities between amino acids. aaHash performs ∼10X faster than generic string hashing algorithms in hashing adjacent *k*-mers.

**Availability and implementation:** aaHash is available online at https://github.com/bcgsc/btllib and is free for academic use.

## Introduction

Analyzing proteins provides opportunities to elucidate more direct insights into the biochemical pathways and functional activities of cells, tissues, and organisms compared to analyzing nucleic acids alone. Conservation at the protein level can reveal important functional and evolutionary insights that may not be immediately apparent when studying sequences at the nucleotide level because of codon degeneracy (Miyata *et al*., 1980). For example, in an assessment of antimicrobial resistance, counting amino acid *k*-mers as opposed to nucleotide *k*-mers was shown to enable higher accuracy and enhanced interpretability of machine learning algorithms (ValizadehAslani *et al*., 2020).

While there are nucleotide-specific hashing algorithms designed for hashing *k*-mers (Mohamadi *et al*., 2016; Kazemi *et al*., 2022), there is no stand-alone and optimized implementation that leverages the characteristics of protein *k-*mers to the best of our knowledge. These nucleotide-specific algorithms first break the sequences into *k*-mers and, typically, map them to 64-bit integers, the largest native type supported by computers. Mapping is achieved by using a hashing algorithm such as ntHash (Mohamadi *et al*., 2016; Kazemi *et al*., 2022), or by encoding nucleic acid characters, Σ = {A, C, G, T|U} using two bits (00, 01, 10, 11) (Simpson *et al*., 2009). Compared to using a hashing algorithm, 2-bit encoding has the advantage of being reversible, but is limited to *k-*mers of length 32bp or shorter using a single 64-bit register. This length limitation is acceptable for certain genomic applications, such as mapping (Li, 2018), polishing (Li *et al*., 2022), or scaffolding (Coombe *et al*., 2021, 2023) but not for other applications, such as de Bruijn graph genome assembly for complex organisms (Jackman *et al*., 2017).

Similarly, amino acids can utilize a 4-or 5-bit encoding, a variant of 2-bit encoding, but this can only capture *k*-mers that are up to 16 amino acid residues in length or shorter using 64 bits, thereby virtually leaving general hashing algorithms as the only viable alternative for hashing longer peptide *k*-mers. Moreover, unlike nucleic acid sequences, where every base substitution is considered equally likely with the use of an identical penalty for any nucleotide mismatch in sequence alignment algorithms (Smith and Waterman, 1981), the interrelationships between amino acids are more complex, necessitating an understanding of molecular structure and biochemical properties, such as hydrophobicity, polarity, and pKa, a measure of the acidity of a molecule (also known as the dissociation constant). Incorporating the relationships between amino acids, for instance, using the BLOSUM62 matrix (Henikoff and Henikoff, 1992), has been shown to improve sensitivity in homology search (Li *et al*., 2009). With high-throughput protein or peptide sequencing platforms on the horizon, researchers will need fast and efficient algorithms tailored for amino acid sequences to rapidly exploit the influx of data and expedite their analyses (Alfaro *et al*., 2021).

In recent years, a number of tools that use amino acid *k-*mers have been developed. KAAmer (Déraspe *et al*., 2022) is a database that utilizes key-value pairs to link amino acid 7-mers, which have been hashed to their corresponding 32-bit hash values by naively combining the hash values of each constituent amino acid with bitwise XOR and shift operations, to the locations of their corresponding contents, i.e., proteins that contain the *k*-mers. Miniprot (Li, 2023), a protein to genome aligner, uses the *k-*mers from a query protein to identify alignment anchors by querying a

6-frame translated genome index using a 4-bit encoding scheme where multiple amino acids are mapped to the same hash value. An optimized amino acid hashing algorithm that does not suffer from the restrictions of 4-or 5-bit encoding could boost the performance of these tools by reducing the processing time and capturing more specific amino acid *k*-mers. Here we present aaHash, a hashing algorithm designed for amino acid sequences that uses multi-level seed tables to represent the biochemical similarities between amino acids.

## Methods

aaHash builds on ntHash (Mohamadi *et al*., 2016; Kazemi *et al*., 2022), a rolling hash algorithm for DNA/RNA sequences, and adapts it for amino acid sequences. Similar to ntHash, aaHash utilizes a seed table to rapidly hash adjacent *k*-mers. This seed table contains 20 arbitrary 64-bit integers indexed using the amino acids alphabet, Σ = {A, C, D, E, F, G, H, I, K, L, M, N, P, Q, R, S, T, V, W, Y}. aaHash also employs pre-built amino acid dimer and trimer seed tables to expedite the calculation of its base hash values (input to recursive hash value calculations) (Supplementary Figure S1).

To leverage the amino acid relationships captured in the BLOSUM62 matrix, we first transformed the matrix such that the positive scores (Supplementary Figure S2), indicating molecular similarity, are concentrated along the diagonal, thereby grouping similar amino acids together (Li *et al*., 2009). We then determined different zones of degeneracy where all the amino acids within the zone should evaluate to the same hash. Using these zones, we created three levels of hashes for aaHash by having seed tables for each level of hash. The first level is a hash function where each amino acid character corresponds to a 64-bit hash value, the default behaviour in generic string hashing. For level 2, we implemented a degenerate hashing scheme where amino acids are grouped when they have positive scores with each other in the BLOSUM62 matrix. Similarly, for level 3, we expanded the definition of degeneracy to include amino acids with nonnegative scores. For hash levels 2 and 3, amino acids in the same degeneracy group will all evaluate to the same hash seed. aaHash also supports using these different levels of hashes together to create a multi-level pattern, mimicking the functionality of spaced seeds (Ma *et al*., 2002).

To evaluate aaHash, we compared its speed, uniformity, and RAM usage against those of CityHash (commit f5dc541), MurmurHash (commit 92cf370), and xxHash (v0.8.1) (Supplementary Table S1). All benchmarking tests were performed using a single thread on a server-class system with 144 Intel(R) Xeon(R) Gold 6254 CPU @ 3.1 GHz with 2.9 TB RAM. aaHash is freely available and implemented within the btllib common code library v1.6.0 (Nikolić *et al*., 2022).

## Results and discussions

To evaluate aaHash’s performance in hashing consecutive amino acid *k*-mers, we hashed 1,000,000 random simulated peptide sequences, each with 250 amino acid residues, using aaHash and three state-of-the-art hashing algorithms, CityHash, MurmurHash, and xxHash (Supplementary Note S1). aaHash is the fastest among all competitors when hashing at least six adjacent *k*-mers, and achieves up to ∼10X speed improvement over CityHash (0.67s vs 7.04s), the second fastest hashing algorithm when hashing 25-mers, i.e. hashing 226 consecutive *k*-mers (Fig. 1a, Supplementary Table S2). aaHash also hashed one billion consecutive 50-mers in 3.19s. 4.99s, and 6.48s to generate 1, 3, and 5 hashes, respectively, again at least an order of magnitude faster than comparators in this use case (Fig. 1b, Supplementary Table S3).

**Fig 1.**
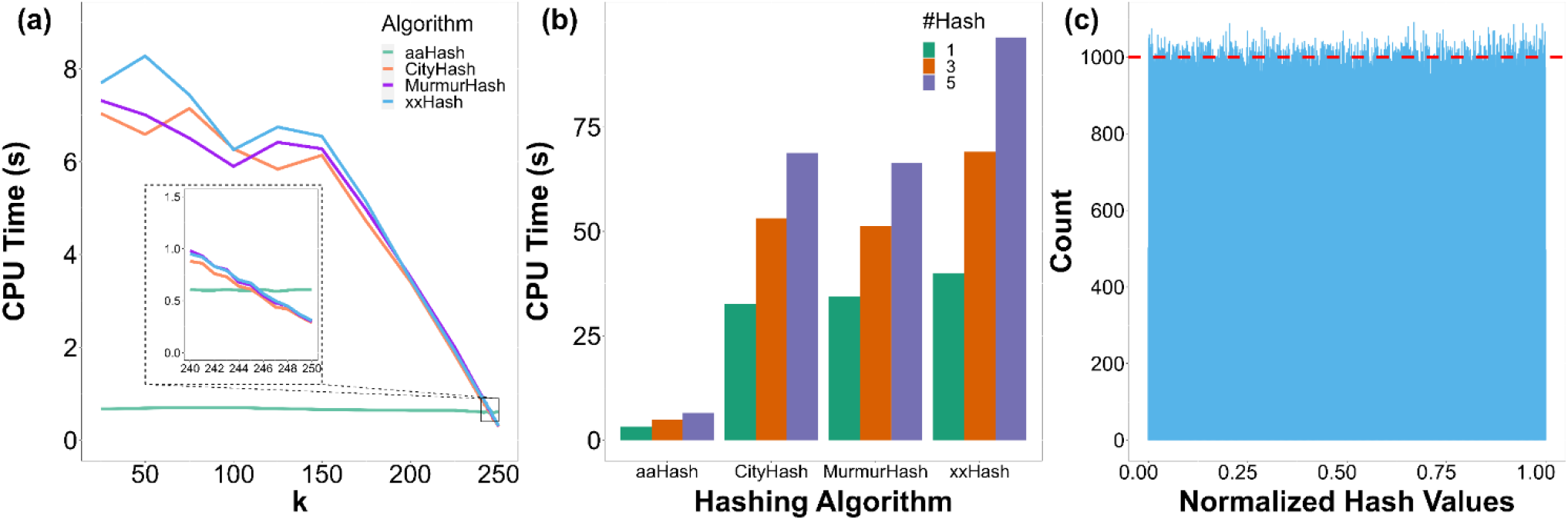
Performance of aaHash. (a) Runtime for hashing 1,000,000 250 amino acids residue long sequences with *k*-mer lengths from 25 to 250. aaHash outperforms all other hashing methods when computing more than five subsequent *k*-mers (i.e. k < 246, see inset). **(b)** Comparing multi-hashing runtime of aaHash versus other state-of-the-art hashing functions for one billion 50-mers. aaHash hashing is ∼10x faster than the closest competitor, CityHash. **(c)** Histogram of 1,000,000 100-mer hashes generated by aaHash from a random amino acid sequence of length 1,000,099. The red dashed line indicates the average number of hashes in a bin (1,000). The hash values were normalized by dividing the hash values by 2^64^ −1, the largest 64-bit integer, and plotted on the histogram with bin size of 1,000. The mean and standard deviation of the bin counts are 1000.0±31.4, demonstrating the uniformity of aaHash.

Next, since uniform distribution of the hashes over the hash space leads to smaller collision probabilities, we evaluated the uniformity of the distribution of aaHash hash values by plotting a histogram of 1,000,000 normalized hash values from a random amino acid sequence (Fig. 1c). Using 1,000 bins, the mean and standard deviation are 1,000.0±31.4, close to the ideal count of 1,000 per bin. We observed similar uniformity and distribution for level 2 and 3 hashes, achieving a mean and standard deviation of 1000.0±31.2 and 1000.0±32.1 (Supplementary Figures S3-4). A Kolmogorov–Smirnov (K-S) test was used to corroborate these observations and showed that the different level hash values aaHash generated are not significantly different from a uniform distribution, an important quality of good hash functions (K-S statistics of 0.0011, 0.0010, and 0.0056 and *P*-values of 0.15, 0.27, and 0.91 for level 1, 2, and 3 hash, respectively) (Chakravarti *et al*., 1967). We also compared the uniformity and distribution of the hash values generated by aaHash and the state-of-the-art hashing algorithms by querying *k*-mers from the UniProt human proteome (Bateman *et al*., 2023) and 1,000,000 randomly simulated 250 amino acid residues long sequences against a Bloom filter (Bloom, 1970), a probabilistic data structure, loaded with *k*-mers from 1,000,000 randomly generated 250 amino acid residues long sequences (Supplementary Note S1, Supplementary Tables S4-9). As the Bloom filter contains only randomly generated *k*-mers, any hits would be considered false positives. We then compared the actual false positive rate with the theoretical false positive rate calculated using the occupancy of the Bloom filter to determine if the aaHash-generated hash values follow a uniform distribution. aaHash achieved false positive rates of 11.9±0.3%, 3.1±0.07, 2.2±0.04% for 1, 3, and 5 hashes, respectively. These do not differ significantly from the theoretical false positive rates of 11.8%, 3.1%, and 2.2% for 1, 3, and 5 hashes, respectively, for both datasets across multiple *k*-mer lengths based on the Student’s paired t-test (t statistics of 1.00, 1.00, and 0.98 and *P*-values of 0.37, 0.36, and 0.36) (Student, 1908). These results are also on par with the other state-of-the-art hashing algorithms and demonstrate that the hash values of aaHash follow a uniform distribution.

Finally, we compared the peak memory usage of each hashing algorithm and note that the peak memory usage does not differ substantially, regardless of *k*-mer length and number of hashes generated (Supplementary Tables S10-11).

Unlike other hashing algorithms, aaHash introduces second and third level hashing, enabling researchers to compare amino acid sequences at the hash level using the BLOSUM62 matrix. BLOSUM62 was chosen because it is the default matrix for BLASTp (Altschul *et al*., 1990), but the concept can be extended to other similarity matrices like PAM matrices (Dayhoff *et al*., 1978). In addition, aaHash supports the integration of various hash levels to produce multi-level patterns. These multi-level patterns mimic the functionality of spaced seeds (a pattern with “care” and “don’t care” positions) (Ma *et al*., 2002), which are typically used for approximate matching in DNA homology searches, but instead of completely ignoring the “don’t care” positions, aaHash will consider the biochemical similarity between the amino acids based on the hash level of each position.

aaHash is a specialized amino acid hashing algorithm that outperforms other state-of-the-art hashing algorithms in speed when hashing consecutive amino acid *k*-mers, a frequently employed operation in bioinformatics. Additionally, the implementation of multi-level hashing has great potential for enabling homology searches between evolutionarily divergent sequences. With its improved speed over other state-of-the art algorithms and homology-oriented features, we expect aaHash to be both beneficial to the scientific community and improve many bioinformatics applications involving amino acid sequence analysis.

## Supplementary Information

Supplementary Information

Supplementary Figs. S1-4, Tables S1, S3-11. Supplementary Table S2

## References

Alfaro, J.A. et al. (2021) The emerging landscape of single-molecule protein sequencing technologies. Nat Methods, 18, 604–617.

Altschul, S.F. et al. (1990) Basic local alignment search tool. J Mol Biol, 215, 403–410.

Bateman, A. et al. (2023) UniProt: the Universal Protein Knowledgebase in 2023. Nucleic Acids Res, 51, D523–D531.

Bloom, B.H. (1970) Space/time trade-offs in hash coding with allowable errors. Commun ACM, 13, 422–426.

Chakravarti, I.M. et al. (1967) Handbook of methods of applied statistics. Wiley Series in Probability and Mathematical Statistics (USA) eng.

Coombe, L. et al. (2021) LongStitch: high-quality genome assembly correction and scaffolding using long reads. BMC Bioinformatics, 22, 534.

Coombe, L. et al. (2023) ntLink: A Toolkit for De Novo Genome Assembly Scaffolding and Mapping Using Long Reads. Curr Protoc, 3.

Dayhoff, M. et al. (1978) A model of evolutionary change in proteins. Atlas of protein sequence and structure, 5, 345–352.

Déraspe, M. et al. (2022) Flexible protein database based on amino acid k-mers. Sci Rep, 12, 9101.

Henikoff, S. and Henikoff, J.G. (1992) Amino acid substitution matrices from protein blocks. Proceedings of the National Academy of Sciences, 89, 10915–10919.

Jackman, S.D. et al. (2017) ABySS 2.0: resource-efficient assembly of large genomes using a Bloom filter. Genome Res, 27, 768–777.

Kazemi, P. et al. (2022) ntHash2: recursive spaced seed hashing for nucleotide sequences. Bioinformatics, btac564.

Li, H. (2018) Minimap2: pairwise alignment for nucleotide sequences. Bioinformatics, 34, 3094–3100.

Li, H. (2023) Protein-to-genome alignment with miniprot. Bioinformatics, 39.

Li, J.X. et al. (2022) ntEdit+Sealer: Efficient Targeted Error Resolution and Automated Finishing of Long-Read Genome Assemblies. Curr Protoc, 2, e442.

Li, W. et al. (2009) Amino Acid Classification and Hash Seeds for Homology Search. In, Rajasekaran, S. (ed), Bioinformatics and Computational Biology. Springer Berlin Heidelberg, Berlin, Heidelberg, pp. 44–51.

Ma, B. et al. (2002) PatternHunter: faster and more sensitive homology search. Bioinformatics, 18, 440–445.

Miyata, T. et al. (1980) Nucleotide sequence divergence and functional constraint in mRNA evolution. Proceedings of the National Academy of Sciences, 77, 7328–7332.

Mohamadi, H. et al. (2016) ntHash: recursive nucleotide hashing. Bioinformatics, 32, 3492–3494.

Nikolić, V. et al. (2022) btllib: A C++ library with Python interface for efficient genomic sequence processing. J Open Source Softw, 7, 4720.

Simpson, J.T. et al. (2009) ABySS: A parallel assembler for short read sequence data. Genome Res, 19, 1117–1123.

Smith, T.F. and Waterman, M.S. (1981) Identification of common molecular subsequences. J Mol Biol, 147, 195–197.

Student (1908) The Probable Error of a Mean. Biometrika, 6, 1.

ValizadehAslani, T. et al. (2020) Amino Acid k-mer Feature Extraction for Quantitative Antimicrobial Resistance (AMR) Prediction by Machine Learning and Model Interpretation for Biological Insights. Biology (Basel), 9, 365.

