## Supplementary Information for "aaHash: recursive amino acid sequence hashing"

**Supplementary Note S1: aaHash benchmarking data**

The dataset we used to generate Supplementary Tables S4-6 are from the UniProt human canonical and isoform proteome (Bateman et al., 2023). The file can be downloaded using this hyperlink: https://doi.org/10.5281/zenodo.7793251.

For Supplementary Figs. 3 and 4, and Supplementary Tables S2 and 3, S7-11, we used random amino acid sequences generated using the btllib library (Nikolić et al., 2022).

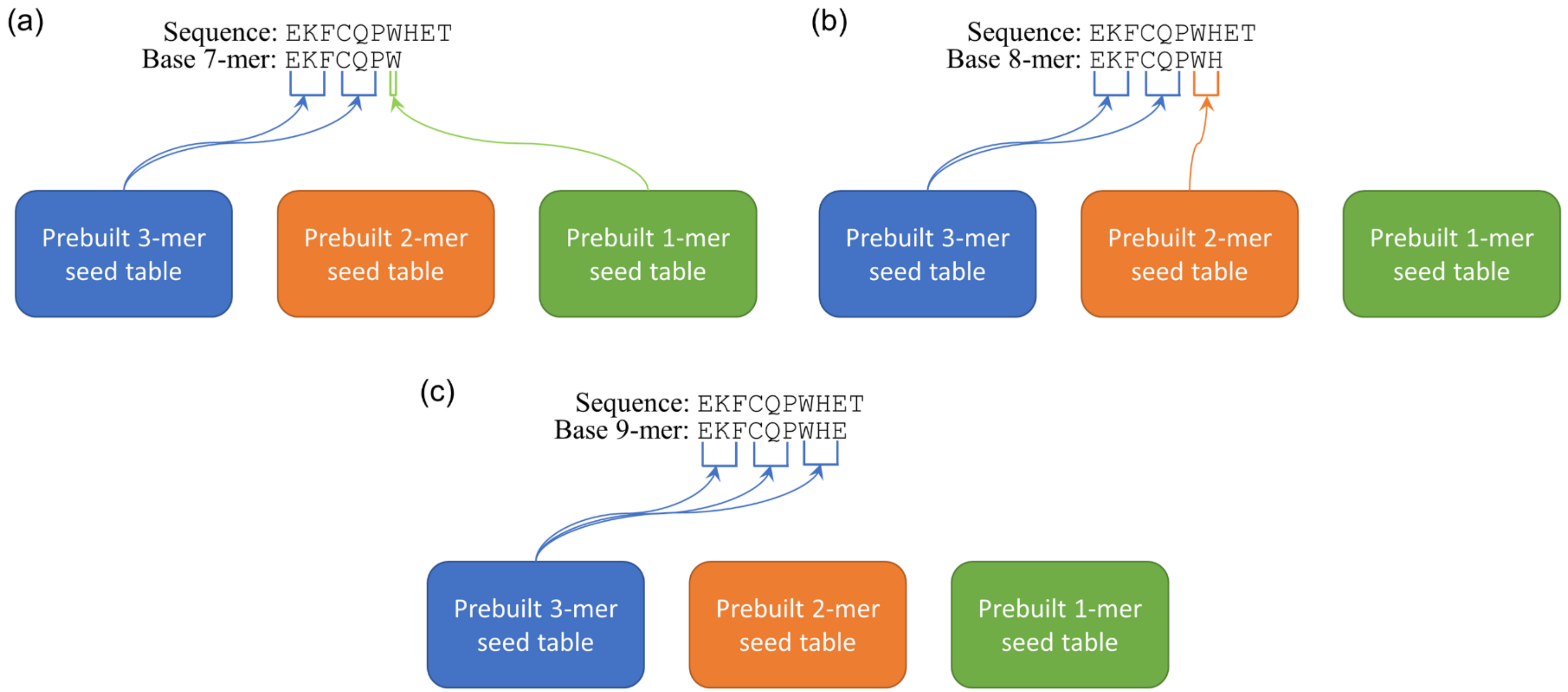

**Supplementary Figure S1. Using pre-built seed tables to compute the base hash value. (a)** Generating base 7-mer hash. **(b)** Generating base 8-mer hash. **(c)** Generating base 9-mer hash. By using prebuilt seed tables, aaHash reduces the number of operations required to create a base hash from $k$, where $k$ is the size of the $k$-mer, to $\lceil k \div 3 \rceil$.

| | C | G | A | T | S | N | D | E | Q | K | R | V | I | L | M | W | F | Y | H | P |
|---|---|---|---|---|---|---|---|---|---|---|---|---|---|---|---|---|---|---|---|---|
| **C** | 9 | -3 | 0 | -1 | -1 | -3 | -3 | -4 | -3 | -3 | -3 | -1 | -1 | -1 | -1 | -2 | -2 | -2 | -3 | -3 |
| **G** | -3 | 6 | 0 | -2 | 0 | 0 | -1 | -2 | -2 | -2 | -2 | -3 | -4 | -4 | -3 | -2 | -3 | -3 | -2 | -2 |
| **A** | 0 | 0 | 4 | 0 | 1 | -2 | -2 | -1 | -1 | -1 | -1 | 0 | -1 | -1 | -1 | -3 | -2 | -2 | -2 | -1 |
| **T** | -1 | -2 | 0 | 5 | 1 | 0 | -1 | -1 | -1 | -1 | -1 | 0 | -1 | -1 | -1 | -2 | -2 | -2 | -2 | -1 |
| **S** | -1 | 0 | 1 | 1 | 4 | 1 | 0 | 0 | 0 | 0 | -1 | -2 | -2 | -2 | -1 | -3 | -2 | -2 | -1 | -1 |
| **N** | -3 | 0 | -2 | 0 | 1 | 6 | 1 | 0 | 0 | 0 | 0 | -3 | -3 | -3 | -2 | -4 | -3 | -2 | 1 | -2 |
| **D** | -3 | -1 | -2 | -1 | 0 | 1 | 6 | 2 | 0 | -1 | -2 | -3 | -3 | -4 | -3 | -4 | -3 | -3 | -1 | -1 |
| **E** | -4 | -2 | -1 | -1 | 0 | 0 | 2 | 5 | 2 | 1 | 0 | -2 | -3 | -3 | -2 | -3 | -3 | -2 | 0 | -1 |
| **Q** | -3 | -2 | -1 | -1 | 0 | 0 | 0 | 2 | 5 | 1 | 1 | -2 | -3 | -2 | 0 | -2 | -3 | -1 | 0 | -1 |
| **K** | -3 | -2 | -1 | -1 | 0 | 0 | -1 | 1 | 1 | 5 | 2 | -2 | -3 | -2 | -1 | -3 | -3 | -2 | -1 | 1 |
| **R** | -3 | -2 | -1 | -1 | -1 | 0 | -2 | 0 | 1 | 2 | 5 | -3 | -3 | -2 | -1 | -3 | -3 | -2 | 0 | -2 |
| **V** | -1 | -3 | 0 | 0 | -2 | -3 | -3 | -2 | -2 | -2 | -3 | 4 | 3 | 1 | 1 | -3 | -1 | -1 | -3 | -2 |
| **I** | -1 | -4 | -1 | -1 | -2 | -3 | -3 | -3 | -3 | -3 | -3 | 3 | 4 | 2 | 1 | -3 | 0 | -1 | -3 | -3 |
| **L** | -1 | -4 | -1 | -1 | -2 | -3 | -4 | -3 | -2 | -2 | -2 | 1 | 2 | 4 | 2 | -2 | 0 | -1 | -3 | -3 |
| **M** | -1 | -3 | -1 | -1 | -1 | -2 | -3 | -2 | 0 | -1 | -1 | 1 | 1 | 2 | 5 | -1 | 0 | -1 | -2 | -2 |
| **W** | -2 | -2 | -3 | -2 | -3 | -4 | -4 | -3 | -2 | -3 | -3 | -3 | -3 | -2 | -1 | 11 | 1 | 2 | -2 | -4 |
| **F** | -2 | -3 | -2 | -2 | -2 | -3 | -3 | -3 | -3 | -3 | -3 | -1 | 0 | 0 | 0 | 1 | 6 | 3 | -1 | -4 |
| **Y** | -2 | -3 | -2 | -2 | -2 | -2 | -3 | -2 | -1 | -2 | -2 | -1 | -1 | -1 | -1 | 2 | 3 | 7 | 2 | -3 |
| **H** | -3 | -2 | -2 | -2 | -1 | 1 | -1 | 0 | 0 | -1 | 0 | -3 | -3 | -3 | -2 | -2 | 1 | 2 | 8 | -2 |
| **P** | -3 | -2 | -1 | -1 | -1 | -2 | -1 | -1 | -1 | -1 | -2 | -2 | -3 | -3 | -2 | -4 | -4 | -3 | -2 | 7 |

**Supplementary Figure S2. Sorted BLOSUM62 matrix**. The BLOSUM (Henikoff & Henikoff, 1992) matrix is sorted such that positive scores, denoting similarity between amino acids, are located on the diagonal. This allows us to cluster similar amino acids together. These clusters are then used to determine which amino acids share the same hash value. The yellow highlighted squares represent which amino acids are hashed to the same value in the level 2 hash (Li et al., 2009). The blue outlined squares represent which amino acids are hashed to the same value in the level 3 hash.

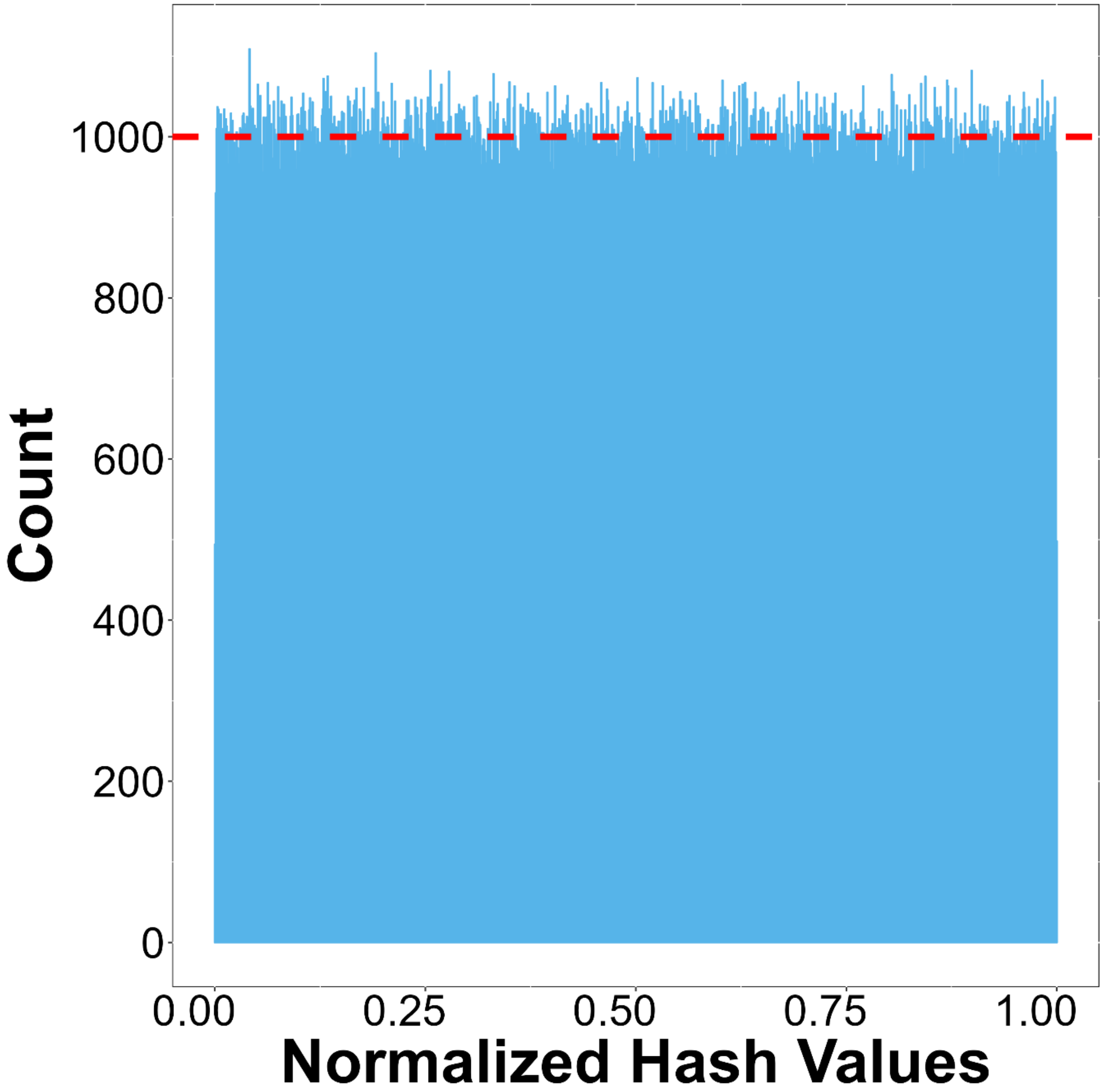

**Supplementary Figure S3. Histogram of 1,000,000 100-mer level 2 hashes generated by aaHash from a random amino acid sequence of length 1,000,099.** The red dashed line indicates the average number of hashes in a bin (1,000). The hash values were normalized by dividing them with $2^{64} - 1$, the largest 64-bit integer, and plotted on the histogram with bin size of 1,000. The mean and standard deviation of the bin counts are 1000.0±31.2, demonstrating the uniformity of aaHash.

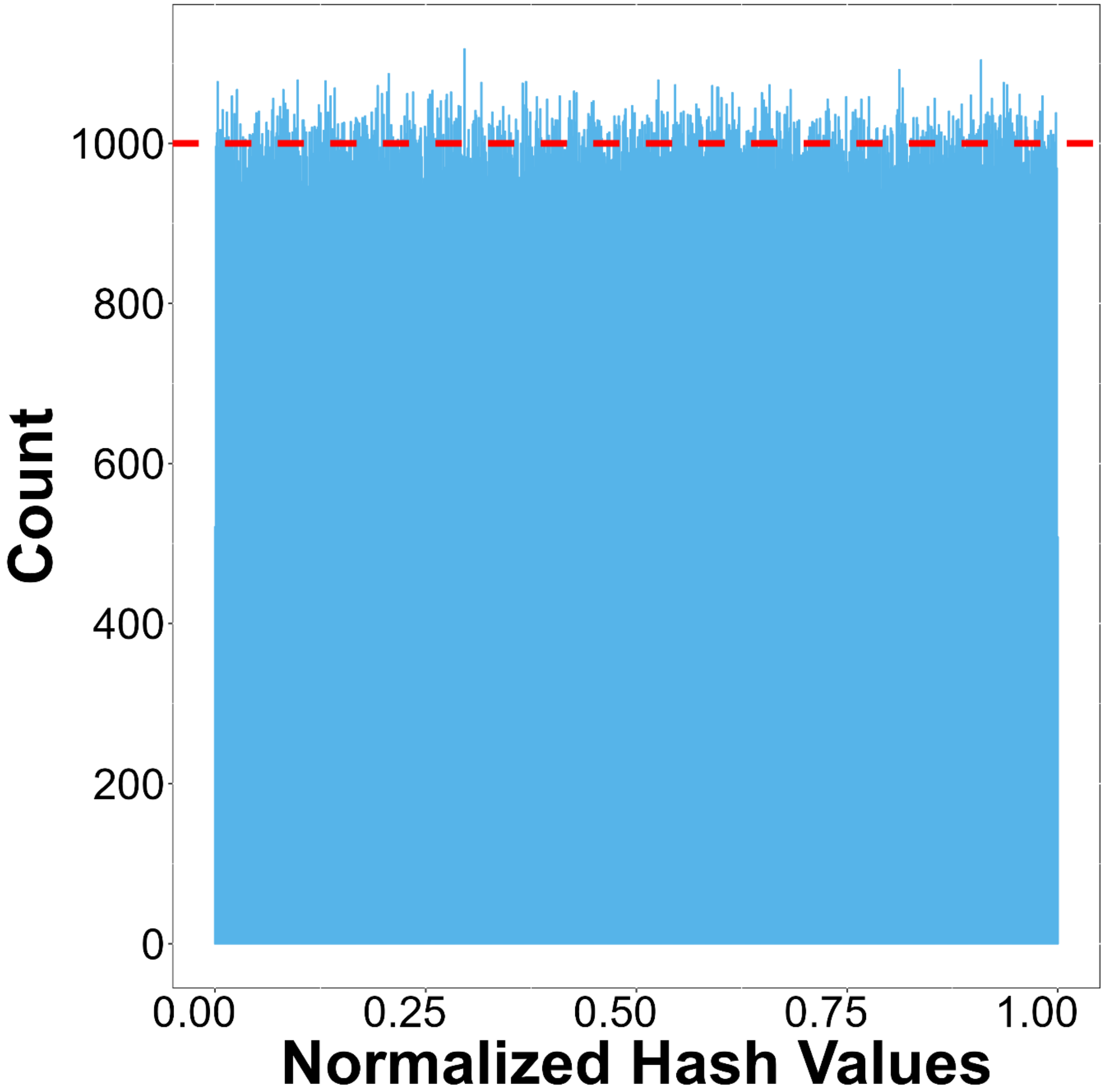

**Supplementary Figure S4. Histogram of 1,000,000 100-mer level 3 hashes generated by aaHash from a random amino acid sequence of length 1,000,099.** The red dashed line indicates the average number of hashes in a bin (1,000). The hash values were normalized by dividing them with $2^{64} - 1$, the largest 64-bit integer, and plotted on the histogram with bin size of 1,000. The mean and standard deviation of the bin counts are 1000.0±32.1, demonstrating the uniformity of aaHash.

**Supplementary Table S1. Hashing algorithms used to benchmark aaHash.** All benchmarking tests were performed using 1 thread on a server-class system with 144 Intel(R) Xeon(R) Gold 6254 CPU @ 3.1 GHz with 2.9 TB RAM. Commit of files are listed when there is no release version of the repository.

| Algorithm | Code repository | Version/commit |
|---|---|---|
| City hash | https://github.com/google/cityhash | f5dc541 |
| Murmur hash | https://github.com/aappleby/smhasher | 92cf370 |
| xxHash | https://github.com/Cyan4973/xxHash | 0.8.1 |

**Supplementary Table S3. Run time evaluation for aaHash and comparators on simulated data generating 1, 3, or 5 hash values for 1,000,000,000 adjacent amino acid 50-mers.**

| Algorithm | Number of hashes | Time (s) |
|---|---|---|
| aaHash | 1 | 3.19 |
| CityHash | | 32.63 |
| MurmurHash | | 34.45 |
| xxHash | | 40.04 |
| aaHash | 3 | 4.99 |
| CityHash | | 53.17 |
| MurmurHash | | 51.27 |
| xxHash | | 69.10 |
| aaHash | 5 | 6.48 |
| CityHash | | 68.77 |
| MurmurHash | | 66.35 |
| xxHash | | 96.47 |

**Supplementary Table S4. Bloom filter evaluation for aaHash and comparators on experimental data using a single hash.** We allocate 8 bits of memory per *k*-mer based on the expected number of *k*-mers in the insertion file and load the Bloom filter with a file containing 1,000,000 randomly generated sequences of length 250 amino acid residues. Then, we query the Bloom filter with the human proteome containing all canonical forms and isoforms with *k*-mers of sizes 50, 150 and 250. The theoretical approximate false positive rate is 11.8%.

| Algorithm | *k*-mer size | Number of Queries | Set bits in BF | False hits | False Positive Rate (%) |
|---|---|---|---|---|---|
| aaHash | 50 | 22,240,331 | 188,943,234 | 2,614,647 | 11.7 |
| CityHash | | | 188,945,163 | 2,612,903 | 11.8 |
| MurmurHash | | | 188,945,552 | 2,614,662 | 11.8 |
| xxHash | | | 18,8947,318 | 2,615,142 | 11.8 |
| aaHash | 150 | 18,005,231 | 94,943,041 | 2,135,966 | 11.9 |
| CityHash | | | 94,940,478 | 2,138,626 | 11.9 |
| MurmurHash | | | 94,938,194 | 2,134,078 | 11.9 |
| xxHash | | | 94,947,078 | 2,138,659 | 11.9 |
| aaHash | 250 | 13,770,131 | 939,841 | 1,725,160 | 12.5 |
| CityHash | | | 939,981 | 1,727,276 | 12.5 |
| MurmurHash | | | 939,866 | 1,724,809 | 12.5 |
| xxHash | | | 940,057 | 1,7281,30 | 12.6 |

**Supplementary Table S5. Bloom filter evaluation for aaHash and comparators on experimental data using 3 hashes.** We allocate 8 bits of memory per *k*-mer based on the expected number of *k*-mers in the insertion file and load the Bloom filter with a file containing 1,000,000 randomly generated sequences of length 250 amino acid residues. Then, we query the Bloom filter with the human proteome containing all canonical forms and isoforms with *k*-mers of sizes 50, 150 and 250. The theoretical approximate false positive rate is 3.1%.

| Algorithm | *k*-mer size | Number of Queries | Set bits in BF | False hits | False Positive Rate (%) |
|---|---|---|---|---|---|
| aaHash | 50 | 22,240,331 | 502,823,792 | 680,002 | 3.1 |
| CityHash | | | 502,848,163 | 680,632 | 3.1 |
| MurmurHash | | | 502,841,165 | 676,344 | 3.0 |
| xxHash | | | 502,840,832 | 680,413 | 3.1 |
| aaHash | 150 | 18,005,231 | 252,675,404 | 555,191 | 3.1 |
| CityHash | | | 252,672,915 | 556,088 | 3.1 |
| MurmurHash | | | 252,675,830 | 553,780 | 3.1 |
| xxHash | | | 252,679,859 | 557,243 | 3.1 |
| aaHash | 250 | 13,770,131 | 2,502,111 | 448,559 | 3.3 |
| CityHash | | | 2,501,524 | 450,201 | 3.3 |
| MurmurHash | | | 2,501,649 | 447,992 | 3.3 |
| xxHash | | | 2,502,508 | 450,084 | 3.3 |

**Supplementary Table S6. Bloom filter evaluation for aaHash and comparators on experimental data using 5 hashes.** We allocate 8 bits of memory per *k*-mer based on the expected number of *k*-mers in the insertion file and load the Bloom filter with a file containing 1,000,000 randomly generated sequences of length 250 amino acid residues. Then, we query the Bloom filter with the human proteome containing all canonical forms and isoforms with *k*-mers of sizes 50, 150 and 250. The theoretical approximate false positive rate is 2.2%.

| Algorithm | *k*-mer size | Number of Queries | Set bits in BF | False hits | False Positive Rate (%) |
|---|---|---|---|---|---|
| aaHash | 50 | 22,240,331 | 747,287,691 | 482,467 | 2.2 |
| CityHash | | | 747,321,437 | 482,516 | 2.2 |
| MurmurHash | | | 747,312,591 | 482,472 | 2.2 |
| xxHash | | | 747,299,027 | 480,699 | 2.2 |
| aaHash | 150 | 18,005,231 | 375,509,789 | 394,163 | 2.2 |
| CityHash | | | 375,509,620 | 395,910 | 2.2 |
| MurmurHash | | | 375,508,139 | 392,860 | 2.2 |
| xxHash | | | 375,509,658 | 393,850 | 2.2 |
| aaHash | 250 | 13,770,131 | 3,717,896 | 318,749 | 2.3 |
| CityHash | | | 3,718,154 | 318,951 | 2.3 |
| MurmurHash | | | 3,717,539 | 317,164 | 2.3 |
| xxHash | | | 3,719,204 | 318,731 | 2.3 |

**Supplementary Table S7. Bloom filter evaluation for aaHash and comparators on simulated data using a single hash.** We allocate 8 bits of memory per *k*-mer based on the expected number of *k*-mers in the insertion file and load the Bloom filter with a file containing 1,000,000 randomly generated sequences of length 250 amino acid residues. Then, we query the Bloom filter with the 1,000,000 simulated sequences of length 250 amino acid residues with *k*-mers of sizes 50, 150 and 250. The theoretical approximate false positive rate is 11.8%.

| Algorithm | *k*-mer size | Number of Queries | Set bits in BF | False hits | False Positive Rate (%) |
|---|---|---|---|---|---|
| aaHash | 50 | 201,000,000 | 188,943,234 | 23,616,753 | 11.8 |
| CityHash | | | 188,945,163 | 23,619,243 | 11.8 |
| MurmurHash | | | 188,945,552 | 23,619,240 | 11.8 |
| xxHash | | | 18,8947,318 | 23,624,417 | 11.8 |
| aaHash | 150 | 101,000,000 | 94,943,041 | 11,867,803 | 11.8 |
| CityHash | | | 94,940,478 | 11,865,281 | 11.8 |
| MurmurHash | | | 94,938,194 | 11,866,971 | 11.8 |
| xxHash | | | 94,947,078 | 11,869,525 | 11.8 |
| aaHash | 250 | 1,000,000 | 939,841 | 117,478 | 11.8 |
| CityHash | | | 939,981 | 117,903 | 11.8 |
| MurmurHash | | | 939,866 | 117,180 | 11.7 |
| xxHash | | | 940,057 | 117,503 | 11.8 |

**Supplementary Table S8. Bloom filter evaluation for aaHash and comparators on simulated data using 3 hashes.** We allocate 8 bits of memory per *k*-mer based on the expected number of *k*-mers in the insertion file and load the Bloom filter with a file containing 1,000,000 randomly generated sequences of length 250 amino acid residues. Then, we query the Bloom filter with the 1,000,000 simulated sequences of length 250 amino acid residues with *k*-mers of sizes 50, 150 and 250. The theoretical approximate false positive rate is 3.1%.

| Algorithm | *k*-mer size | Number of Queries | Set bits in BF | False hits | False Positive Rate (%) |
|---|---|---|---|---|---|
| aaHash | 50 | 201,000,000 | 502,823,792 | 6,146,485 | 3.1 |
| CityHash | | | 502,848,163 | 6,148,676 | 3.1 |
| MurmurHash | | | 502,841,165 | 6,144,972 | 3.1 |
| xxHash | | | 502,840,832 | 6,146,218 | 3.1 |
| aaHash | 150 | 101,000,000 | 252,675,404 | 3,088,838 | 3.1 |
| CityHash | | | 252,672,915 | 3,086,423 | 3.1 |
| MurmurHash | | | 252,675,830 | 3,088,088 | 3.1 |
| xxHash | | | 252,679,859 | 3,088,804 | 3.1 |
| aaHash | 250 | 1,000,000 | 2,502,111 | 30,466 | 3.1 |
| CityHash | | | 2,501,524 | 30,812 | 3.1 |
| MurmurHash | | | 2,501,649 | 30,450 | 3.1 |
| xxHash | | | 2,502,508 | 30,483 | 3.1 |

**Supplementary Table S9. Bloom filter evaluation for aaHash and comparators on simulated data using 5 hashes.** We allocate 8 bits of memory per *k*-mer based on the expected number of *k*-mers in the insertion file and load the Bloom filter with a file containing 1,000,000 randomly generated sequences of length 250 amino acid residues. Then, we query the Bloom filter with the 1,000,000 simulated sequences of length 250 amino acid residues with *k*-mers of sizes 50, 150 and 250. The theoretical approximate false positive rate is 2.2%.

| Algorithm | *k*-mer size | Number of Queries | Set bits in BF | False hits | False Positive Rate (%) |
|---|---|---|---|---|---|
| aaHash | 50 | 201,000,000 | 747,287,691 | 4,355,434 | 2.2 |
| CityHash | | | 747,321,437 | 4,359,290 | 2.2 |
| MurmurHash | | | 747,312,591 | 4,356,975 | 2.2 |
| xxHash | | | 747,299,027 | 4,355,402 | 2.2 |
| aaHash | 150 | 101,000,000 | 375,509,789 | 2,188,433 | 2.2 |
| CityHash | | | 375,509,620 | 2,187,385 | 2.2 |
| MurmurHash | | | 375,508,139 | 2,189,370 | 2.2 |
| xxHash | | | 375,509,658 | 2,188,529 | 2.2 |
| aaHash | 250 | 1,000,000 | 3,717,896 | 21,807 | 2.2 |
| CityHash | | | 3,718,154 | 21,536 | 2.2 |
| MurmurHash | | | 3,717,539 | 21,639 | 2.2 |
| xxHash | | | 3,719,204 | 21,401 | 2.1 |

**Supplementary Table S10. Benchmarking script peak memory usage of each hashing algorithm generating 1 hash value for each *k*-mer with different *k*-mer lengths from 1,000,000 simulated sequences of length 250 amino acid residues.**

| Algorithm | *k*-mer size | Peak memory usage (MB) |
|---|---|---|
| aaHash | 50 | 2.1 |
| CityHash | | 2.1 |
| MurmurHash | | 2.1 |
| xxHash | | 2.1 |
| aaHash | 150 | 2.1 |
| CityHash | | 2.1 |
| MurmurHash | | 2.1 |
| xxHash | | 2.1 |
| aaHash | 250 | 2.1 |
| CityHash | | 2.1 |
| MurmurHash | | 2.1 |
| xxHash | | 2.1 |

**Supplementary Table S11. Benchmarking script peak memory usage of each hashing algorithm generating 1, 3, or 5 hash values for each *k*-mer with *k*=50 from 1,000,000 simulated sequences of length 250 amino acid residues.**

| Algorithm | Number of Hashes | Peak memory usage (MB) |
|---|---|---|
| aaHash | 1 | 2.1 |
| CityHash | | 2.1 |
| MurmurHash | | 2.1 |
| xxHash | | 2.1 |
| aaHash | 3 | 2.1 |
| CityHash | | 2.1 |
| MurmurHash | | 2.1 |
| xxHash | | 2.1 |
| aaHash | 5 | 2.1 |
| CityHash | | 2.1 |
| MurmurHash | | 2.1 |
| xxHash | | 2.1 |

**Supplementary References**

Bateman, A., Martin, M.-J., Orchard, S., Magrane, M., Ahmad, S., Alpi, E., Bowler-Barnett, E. H., Britto, R., Bye-A-Jee, H., Cukura, A., Denny, P., Dogan, T., Ebenezer, T., Fan, J., Garmiri, P., da Costa Gonzales, L. J., Hatton-Ellis, E., Hussein, A., Ignatchenko, A., … Zhang, J. (2023). UniProt: the Universal Protein Knowledgebase in 2023. *Nucleic Acids Research*, *51*(D1), D523–D531. https://doi.org/10.1093/nar/gkac1052

Henikoff, S., & Henikoff, J. G. (1992). Amino acid substitution matrices from protein blocks. *Proceedings of the National Academy of Sciences*, *89*(22), 10915–10919. https://doi.org/10.1073/pnas.89.22.10915

Li, W., Ma, B., & Zhang, K. (2009). Amino Acid Classification and Hash Seeds for Homology Search. In S. Rajasekaran (Ed.), *Bioinformatics and Computational Biology* (pp. 44–51). Springer Berlin Heidelberg.

Nikolić, V., Kazemi, P., Coombe, L., Wong, J., Afshinfard, A., Chu, J., Warren, R. L., & Birol, I. (2022). btllib: A C++ library with Python interface for efficient genomic sequence processing. *Journal of Open Source Software*, *7*(79), 4720. https://doi.org/10.21105/joss.04720
